# The response of multi-disease and insect-resistant tomato lines to the accumulation of TYLCV, a whitefly-transmitted virus

**DOI:** 10.1101/2024.09.21.614278

**Authors:** Shruthi Shimoga Prabhakar, Yun-che Hsu, Joyce Yen, Hsiu-yi Chou, Mei-ying Lin, M. Shanthi Priya, Stephen Othim, Srinivasan Ramasamy, Assaf Eybishitz

## Abstract

Tomato (Solanum lycopersicum) is a major vegetable crop grown worldwide for its culinary versatility and nutritional richness. The whitefly-transmitted Tomato yellow leaf curl virus (TYLCV) poses a significant threat to its cultivation. The management strategies for this disease include controlling the virus and/or the vector. The multi-disease and insect-resistant lines developed by the World Vegetable Centre (WorldVeg) contain Ty-1/Ty-3 genes for virus resistance and WF2-10 and WF3-09 genes for whitefly resistance. This study evaluates the efficacy of suppressing virus accumulation in multi-disease and insect-resistant tomato lines compared to Ty-resistant or whitefly-resistant lines and a susceptible check. Preference bioassays, controlled inoculation with viruliferous whiteflies, and acylsugar quantification revealed that multi-disease and insect-resistant lines, developed at WorldVeg, had significantly higher acylsugar concentration and were less preferred by the whitefly adults for settling and whiteflies had high adult mortality. The multi-disease and insect-resistant lines also showed less severe disease symptoms and reduced virus accumulation over time when compared to Ty-resistant, whitefly-resistant, and susceptible check. The findings reveal that the multi-disease and insect-resistant lines are superior in mitigating the threat posed by TYLCV compared to TYLCV-resistant lines. These results underscore the potential of combined virus and vector resistance in tomatoes as a key element of Integrated Pest Management strategies against whitefly-transmitted TYLCV, offering a sustainable solution for safeguarding tomato production.

## Introduction

Tomatoes are one of the most widely consumed vegetables worldwide, for fresh consumption and processing. Tomatoes are highly valued for their nutritional richness and culinary versatility. Globally, tomato cultivation spans an area of approximately 5 million hectares, yielding a production of nearly 186.82 million tonnes with an average productivity of 36.97 tonnes per hectare [1].

Biotic stress for plants includes pests and diseases, which account for significant crop losses threatening crop yield and productivity [2]. The whitefly *Bemisia tabaci* Gennadius (Hemiptera: Aleyrodidae) feeds on many crops, including tomatoes. Its feeding causes damage to the phloem and physiological disorders such as irregular fruit ripening in tomatoes, which reduces fruit quality and increases the number of unmarketable fruit [3]. Whitefly also supports the growth of sooty mold fungus by secreting honeydew. However, its role as a plant virus vector causes the most serious damage [4]. Virus species belonging to five different genera (*Begomovirus, Crinivirus, Closterovirus, Ipomovirus,* and *Carlavirus*) are transmitted by whiteflies [5]. Among the viruses that can infect tomatoes, tomato yellow leaf curl virus (TYLCV) threatens tomato production and currently ranks third after tobacco mosaic virus and tomato spotted wilt virus on the list of the most important plant viruses worldwide [6]. TYLCV is mainly restricted to the tomato phloem tissues and is transmitted in a persistent circulative manner [7]. Tomato Yellow Leaf Curl Disease (TYLCD) symptoms include leaf yellowing, curling, stunting, and, in severe cases, the flowers and fruits are abscised, followed by cessation of plant growth [8]. Early tomato infection can cause up to 100 % yield loss [9].

For profitable tomato production, TYLCD management is very important. Possible measures to control the spread and effects of TYLCV are breeding cultivars resistant to the virus and resistant to the vector. Resistance to TYLCD was found in several wild relatives of tomato, from which six TYLCV resistance genes (*Ty-1* to *Ty-6*) have been identified. *Ty-1* and *Ty-3* are the primary resistance genes widely used in tomato breeding programs. *Ty-2* is also used in breeding, either alone or in combination with other Ty-genes [8]. Ty genes have been successful in generating TYLCV-resistant commercial tomato cultivars, but generally only reduce symptom severity and the resistance achieved has never been 100% [10]. Plant co-infection by multiple Begomovirus strains offers opportunities to recombine and evolve new virulent, resistance-breaking forms of the virus. One more disadvantage of relying on plants with only Ty genes is that they constitute virus sources for susceptible genotypes [11].

While chemical vector control is ineffective, host plant resistance to insect vectors is suitable for managing circulative vector-borne virus diseases. Durable whitefly resistance, especially in the field, is more likely if tomato cultivars mount resistance based on a combination of antixenosis and antibiosis factors, thus forcing whiteflies to surmount a wide range of plant defenses [12]. Physically, plants deploy trichome and acyl sugar-based strategies to restrain whiteflies from feeding. There are two groups of trichomes: glandular trichomes (Type I, IV, VI, and VII trichomes), which have “heads” containing various sticky and/or toxic exudates and secrete acyl sugars and non-glandular trichomes (Type II, III and V trichomes) that do not secrete acyl sugars. Acyl sugars confer resistance to a wide range of important pests [12–14]. Natural resistance to multiple pests based on secretions by the different types of glandular trichomes present on stems and leaves of tomato and its wild relatives has been described [15]. In most cases, feeding deterrence is the mode of resistance, as is oviposition deterrence for some pests. The capacity to repel and avoid whitefly landing, probing, and feeding is important to thwart whiteflies as vectors of plant viruses, especially begomoviruses.

However, vector resistance alone would only lead to reduced virus spread while the virus still can accumulate in plants without any virus resistance mechanism to slow down its replication. Thus, Integrated Pest Management (IPM) is required for a holistic approach to the management of TYLCD. IPM can include along with other measures, host plant resistance to both the whitefly and the virus. Dual resistance in tomato cultivars would be valuable in repelling whiteflies and inhibiting virus replication in the plant, thus helping preserve the durability of virus resistance genes and possibly contributing to slowing Begomovirus evolution.

WorldVeg has developed BC_4_F_5_ breeding lines that are multi-disease and insect-resistant. The accession *VI007099* of *S. galapagense*, a close wild relative to the cultivated tomato, is resistant to the whitefly because it possesses type IV glandular trichomes that produce acylsucroses [12]. This resistance trait was introgressed into the multi-disease resistant elite line *CLN3682C*, and after four steps of recurrent backcrossing with selection for two whitefly resistance markers and acyl sugar quantification breeding lines *AVTO2428*, *AVTO2432*, *AVTO2437,* and *AVTO2436* were selected from *CLN4636BC4F5*. These lines contain *Ty-1/Ty-3* genes for virus resistance *WF2-10* and *WF3-09* genes for whitefly resistance.

So far, little has been done on the response of insect-resistant lines to the accumulation and spread of the TYLCV virus [16–19]. The reaction of dual-resistant lines to the virus accumulation was not studied extensively. The question arises as to whether dual resistance against the insect vector and the virus limits Begomovirus accumulation, because vector resistance can, in some cases, increase virus transmission [20,21]. Vector-resistant cultivars, however, have helped to reduce the spread of other plant viruses [22]. The specific virus–vector interactions that determine Begomovirus transmission are complex, involving the virus and vector, the host plant, and the environment.

Therefore, this study was conducted with the main objective of comparing the TYLCV virus accumulation in plants combining virus and insect resistance with plants either resistant to the insect or the virus alone, and susceptible plants.

## Materials and methods

The experiment was performed under greenhouse conditions (26.7 ± 0.9 ° C temperature; 69.11 ± 1.05% relative humidity) at the WorldVeg in Tainan, Taiwan, from 23 February to 22 May, 2024.

### Tomato plants, virus isolate, and whitefly population

Four multi-disease and insect-resistant lines, *AVTO2428*, *AVTO2432*, *AVTO2437* and *AVTO2436*; one TYLCV-resistant line, *AVTO2445* (stable and advanced line of the recurrent female parent of the multi-disease and virus resistant lines), one whitefly resistant check *AVTO2446* (*S. galapagense*) and one susceptible check *AVTO9304* were used. The four multi-disease and insect-resistant lines are BC_4_F_5_ lines developed by transferring insect resistance from *S. galapagense* accession into an elite multiple disease-resistant (including TYLCV resistance) line. The characteristics of the lines used in the study are summarised in Table 1.

**Table 1.** Characteristics of lines used in the study.

| Genotype | Pedigree | Characteristic | Ty1,3 | WF2-10 | WF3-09 |
| --- | --- | --- | --- | --- | --- |
| **AVTO2428** | CLN4636BC4F2-31I03-1(a)-BK(e) | Multi-disease and insect-resistance line | + | + | + |
| **AVTO2432** | CLN4636BC4F2-31I03-1(d)-BK(e) | Multi-disease and insect-resistance line | + | + | + |
| **AVTO2436** | CLN4636BC4F2-31I03-1(c)-1(d) | Multi-disease and insect-resistance line | + | + | + |
| **AVTO2437** | CLN4636BC4F2-31I03-1(c)-4(a) | Multi-disease<br>and insect-resistance line | + | + | + |
| **AVTO2445** | CLN4396F1-2-18Z85-23-16-9-6 | TYLCV-resistant line | + | — | — |
| **AVTO2446** | VI063177-10<br>(BL1921) | Whitefly-resistant line | — | + | + |
| **AVTO9304** | CL5915-93D4-1-0-3 | Susceptible<br>check | — | — | — |

\*+ Presence of resistance allele ; — Absence of resistance allele.

Tomato plants infected with the TYLCV Thailand strain (TYLCTHV) were obtained from the virology department of WorldVeg.

Healthy *B. tabaci* individuals were obtained from the WorldVeg virology department and reared on cabbage plants in insect cages in an insect-proof plastic house. Viruliferous whiteflies were obtained by releasing healthy adults on TYLCTHV-infected tomato plants and allowing them to feed on them for four days.

### Whitefly preference and TYLCV control inoculation

A choice bioassay was conducted to study the whitefly preference. The 28-day-old seedlings of three plants of each of the 7 genotypes were placed randomly in a circle within insect cages. 70 *B. tabaci* adult mating pairs were released into the cage resulting in 10 mating pairs/genotype. The viruliferous whiteflies were released in the center of the circle. The inoculation access period (IAP) was 8 days (192 hr). After the IAP, the whitefly adults were counted on the whole plants, and then the plants were treated with insecticide.

### TYLCV virus accumulation

TYLCTHV was detected by Polymerase Chain Reaction (PCR). The DNA extraction procedure was performed according to Tanksley Lab [23], modified by WorldVeg. The genomic DNA was quantified through nanodrop (Thermo Scientific NanoDrop spectrometer) and then diluted to 10 ng/μl. The diluted sample was subjected to PCR. Each plant was sampled and tested for the presence and amount of TYLCV in the plants (3-, 5- and 20-days post-inoculation) by comparing with the standard curve developed using serially diluted plasmid DNA at different cycles as shown in Fig 1.

**Fig 1.**
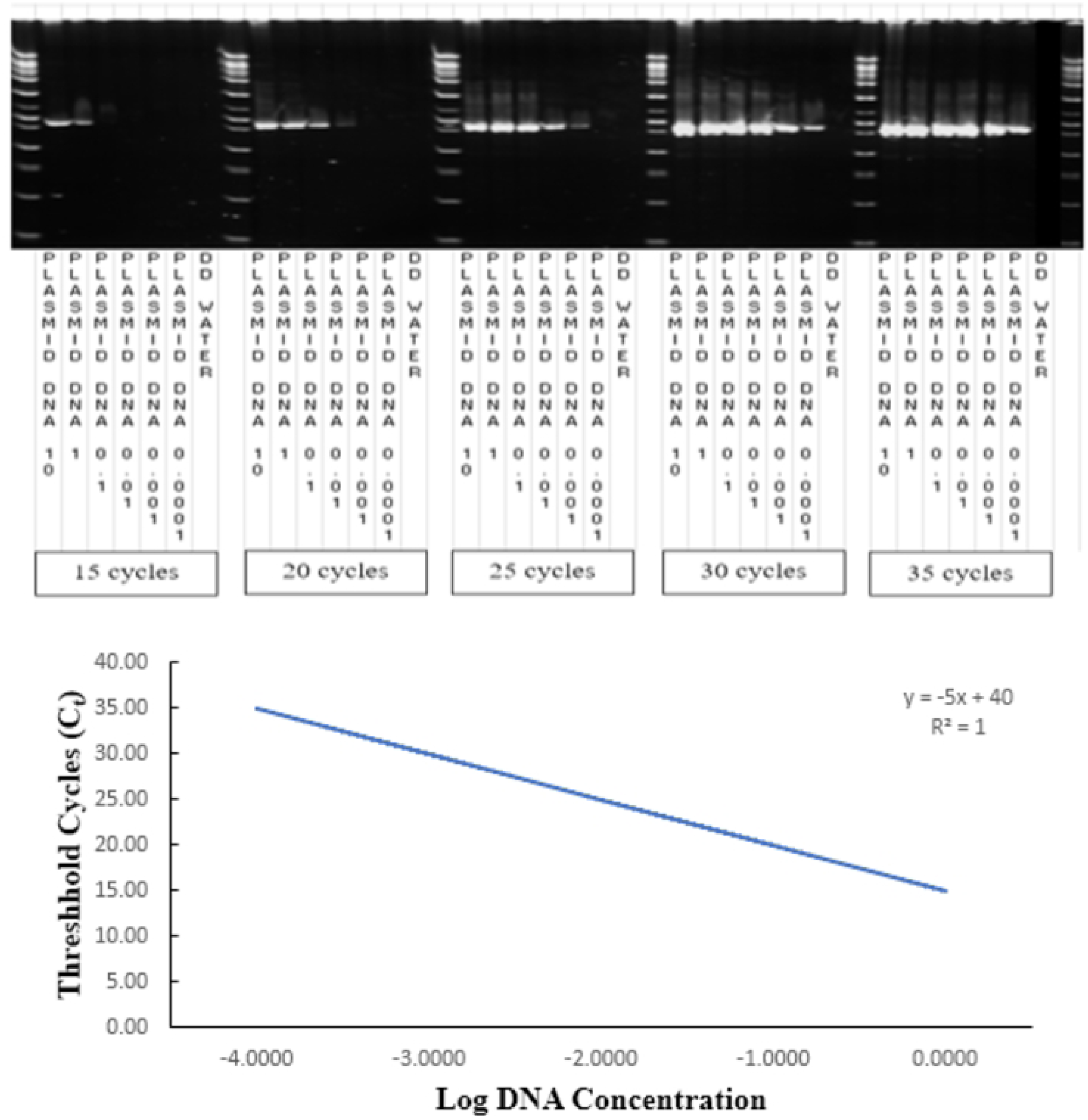
Gel electrophoresis of PCR products of serially diluted plasmid DNA at different cycles and standard curve :threshold cycles(C_t_) vs log DNA concentration (y = - 5x + 40).

To quantify the virus in that sample, the PCR was done at a different number of cycles (15, 20, 25, and 30 cycles) with serially diluted plasmid DNA.

### Disease scoring

The plants were phenotypically evaluated and scored based on the severity of the disease symptoms using a 1-6 scale as described in Table 2. The phenotyping scale is also shown in Fig 2.

**Table 2.** Phenotyping scale for Tomato yellow leaf curl disease (TYLCD).

| Score | Reaction | Description |
| --- | --- | --- |
| 1 | None<br>(Healthy) | No visible symptoms on leaves |
| **2** | Very mild | Slight yellowing, mosaic on youngest (top) leaves, no leaf curl |
| **3** | Mild | Mild yellowing, mosaic and/or slight leaf curling on youngest (top) leaves |
| **4** | Moderate | Moderate yellowing and/or moderate leaf curling on the youngest (top) leaves |
| **5** | Severe | Severe yellowing and blistering and/or severe leaf curling plus some leaf size reduction on the youngest (top) leaves of the main stem and/or on at least one branch |
| **6** | Very severe | Severe yellowing, blistering and/or very severe leaf curling, leaf deformation, leaf size reduction, and stunting (internode-shortening) to the top of the main stem and/or at least one branch |

**Fig 2.**
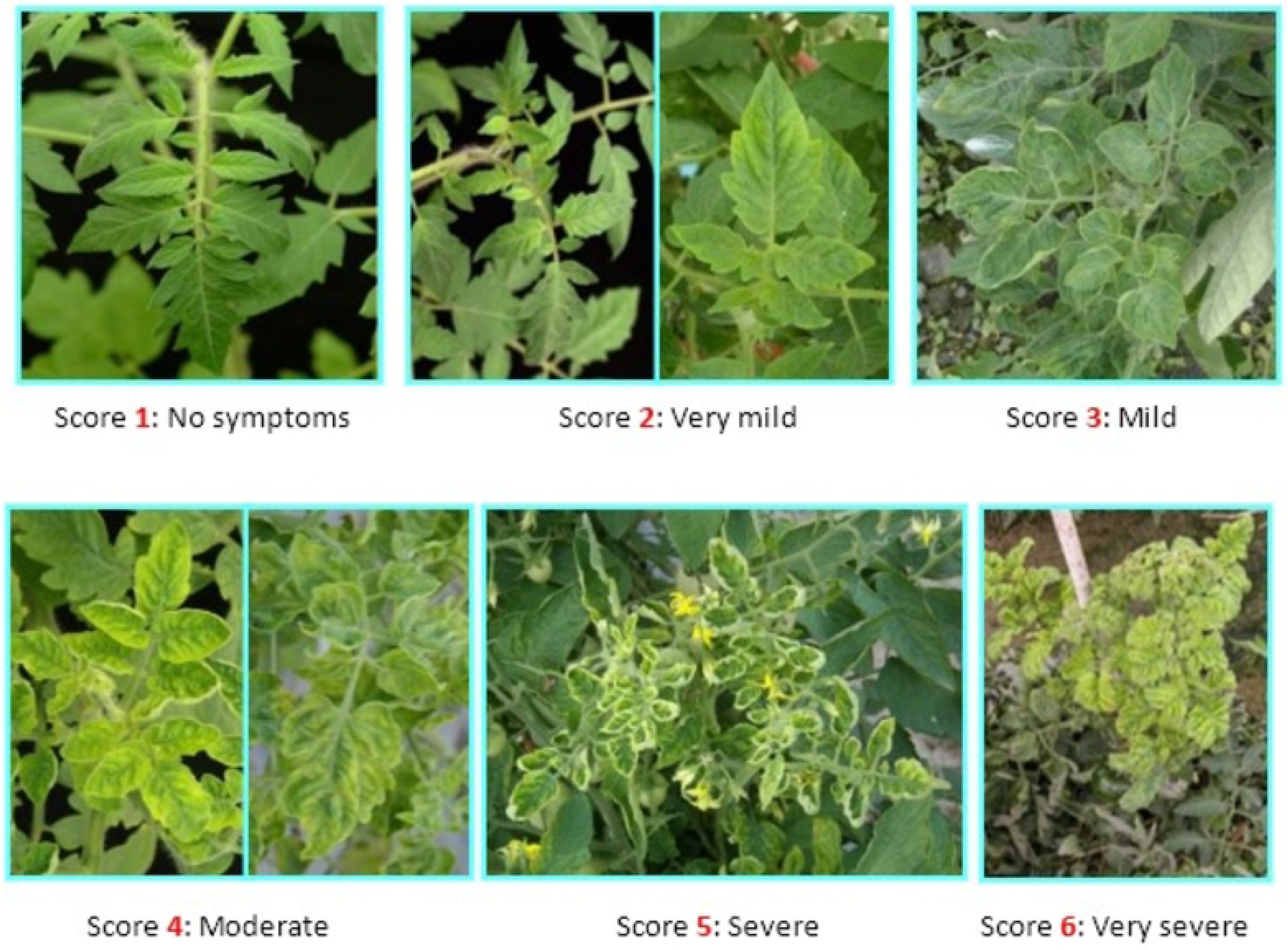
Phenotyping scale-Disease Scoring.

### Acylsugar assay

Standard PGO (Peroxidase/glucose oxidase) based acylsugar assay was performed following the methodology described by [24].

### Statistical analysis

Data were subjected to statistical analysis using SPSS for Windows, version 25.0 (SPSS Inc., Chicago, IL).The data was checked for normality and variance homogeneity using Shapiro-Wilk and Levene’s tests, respectively. A Sqrt(x) transformation was applied to normalize the adult whitefly data, whereas an arcsine(x) transformation was applied to adult mortality (%). Analysis of variance (ANOVA) was performed, and the means were compared through Tukey’s HSD test (p<0.05). The Kruskal–Wallis test was performed for acylsugar content, virus accumulation, and disease severity index, followed by Dunn’s test (p <0.05) for pairwise comparison wherever a statistically significant difference was found.

## Results

A series of controlled greenhouse experiments were conducted to assess the response of multi-disease insect-resistant tomato lines to the spread and accumulation of TYLCV. The results are summarised below.

### Acyl sugar content in leaves

The total acylsugar concentration in different genotypes is presented in Fig 3. The total acylsugar concentration was lower in *AVTO2445* and *AVTO9304*. It was highest in *AVTO2428,* followed by *AVTO2432*, *AVTO2437*, *AVTO2436,* and *AVTO2446*. A Kruskal-Wallis test indicated that there was significant difference in acylsugar content across the 7 genotypes (df=6, N = 21; test statistic χ2= 14.892 p=0.021). Post-hoc comparisons using Dunn’s test indicated that the acylsugar concentration of *AVTO2445* and *AVTO9304* were not significantly different from each other but were significantly different from the multi-disease and insect-resistant lines viz., *AVTO2428*, *AVTO2432,* and *AVTO2437*.

**Fig 3.**
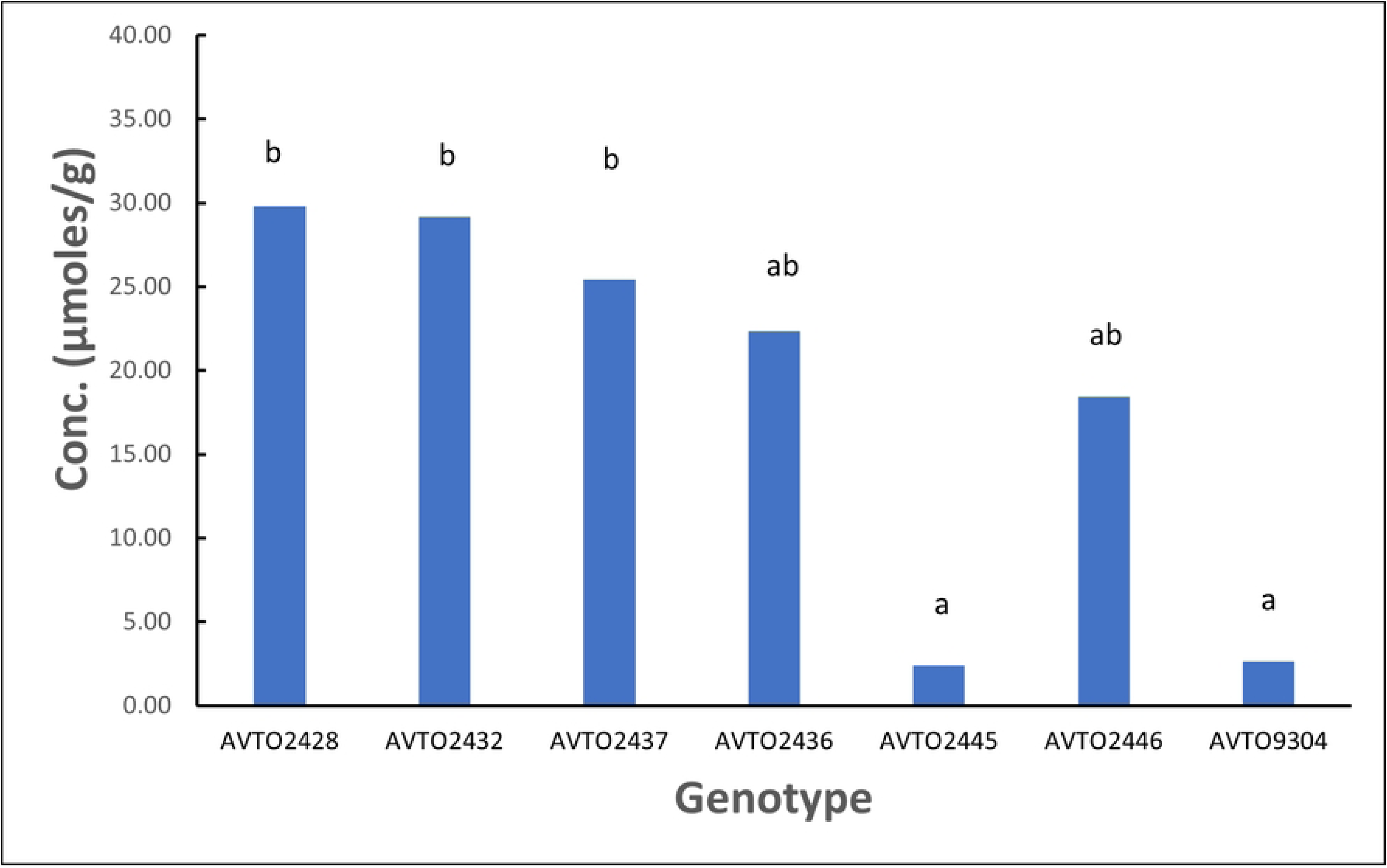
Total acylsugar concentration (μmoles/g). Different letters indicate statistical differences according to Dunn’s tests (P < 0.05).

### Whitefly preference assay

The genotype *AVTO2446* attracted the least whitefly adults in the choice assay as depicted in Fig 4. A one-way ANOVA was conducted to compare the effect of the genotypes on number of adult whitefly/plant in choice assay. A statistically significant difference between the genotypes was found with p <0.05 [F(6, 14) = 3.745, p = 0.020]. Post hoc comparisons using Tukey’s HSD test indicated that the mean score for *AVTO2446* (M = 0.74, SD = 0.65) significantly differed from *AVTO9304* (M = 2.55, SD = 0.48). However, *AVTO2428*, *AVTO2432*, *AVTO2437*, *AVTO2436,* and *AVTO2445* did not significantly differ from each other or from *AVTO2446* or *AVTO9304*. Adult mortality (%) was also high in the multi-disease and insect-resistant lines (Fig 5).

**Fig 4.**
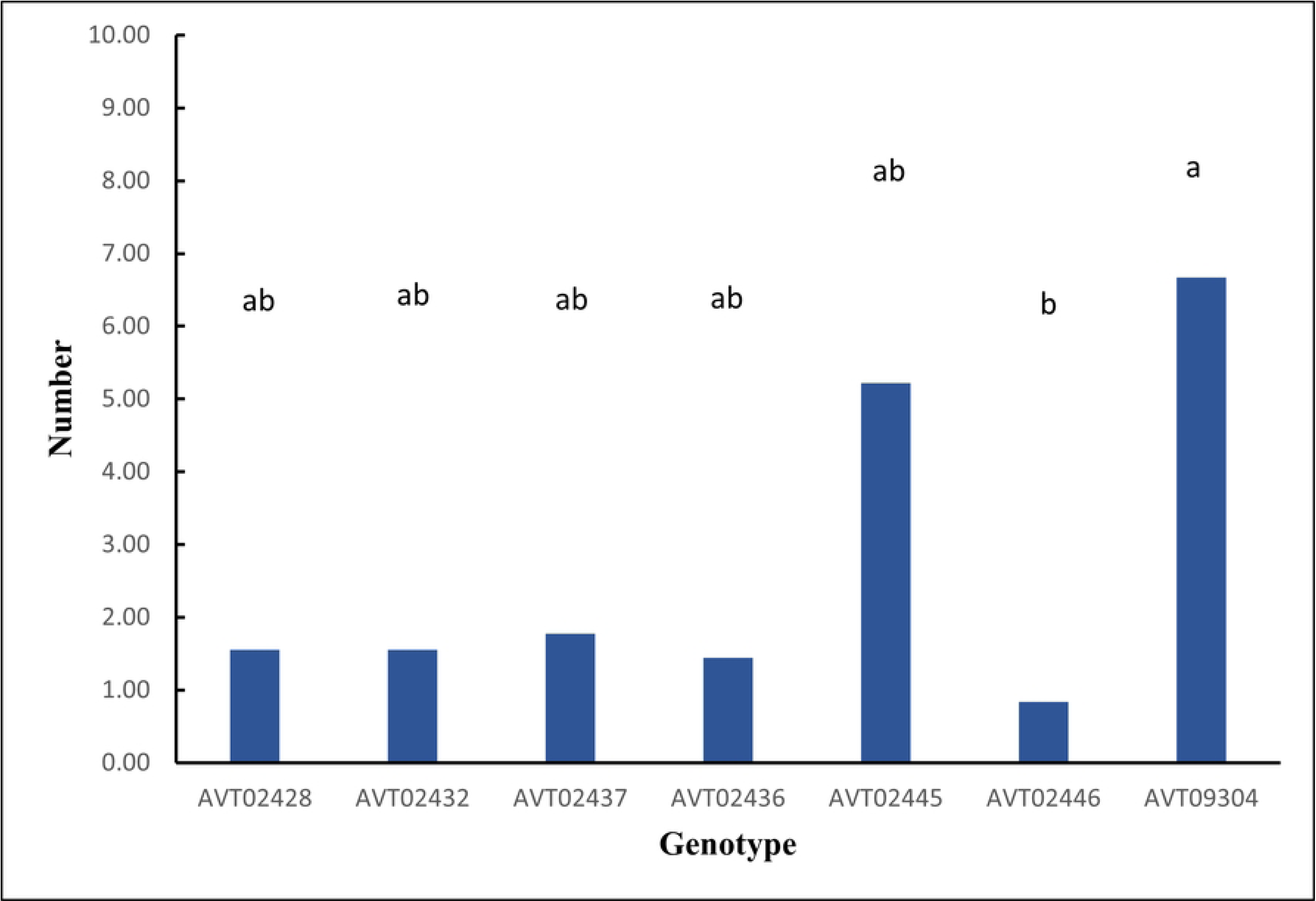
Number of adult whiteflies/plant. Different letters indicate statistical differences according to Tukey’s HSD test (P < 0.05).

**Fig 5.**
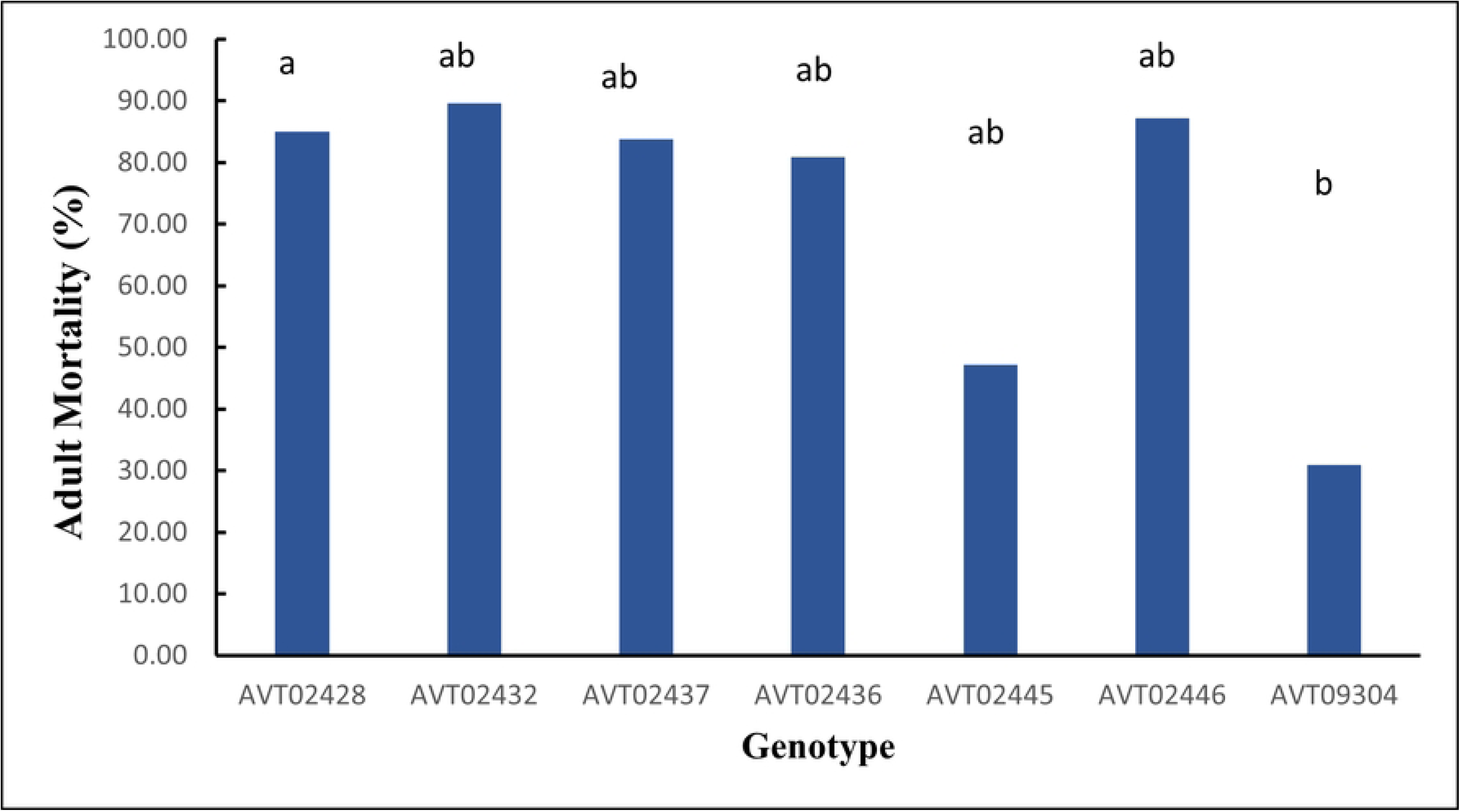
Adult mortality (%). Different letters indicate statistical differences according to Tukey’s HSD test (P < 0.05).

### TYLCV accumulation

A Kruskal-Wallis test was conducted, followed by post hoc comparisons using Dunn’s test wherever a significant difference was found. Table 3 and Fig 6 provides a summary of the viral load on various days post-inoculation. The multi-disease and insect-resistant plants consistently displayed lower viral loads than *AVTO2445*, *AVTO2446,* and *AVTO9304*. At 20 days post-inoculation (dpi), the genotype *AVTO2432* had significantly lower virus amounts compared to *AVTO2445*, *AVTO2446,* and *AVTO9304*.

**Table 3.**
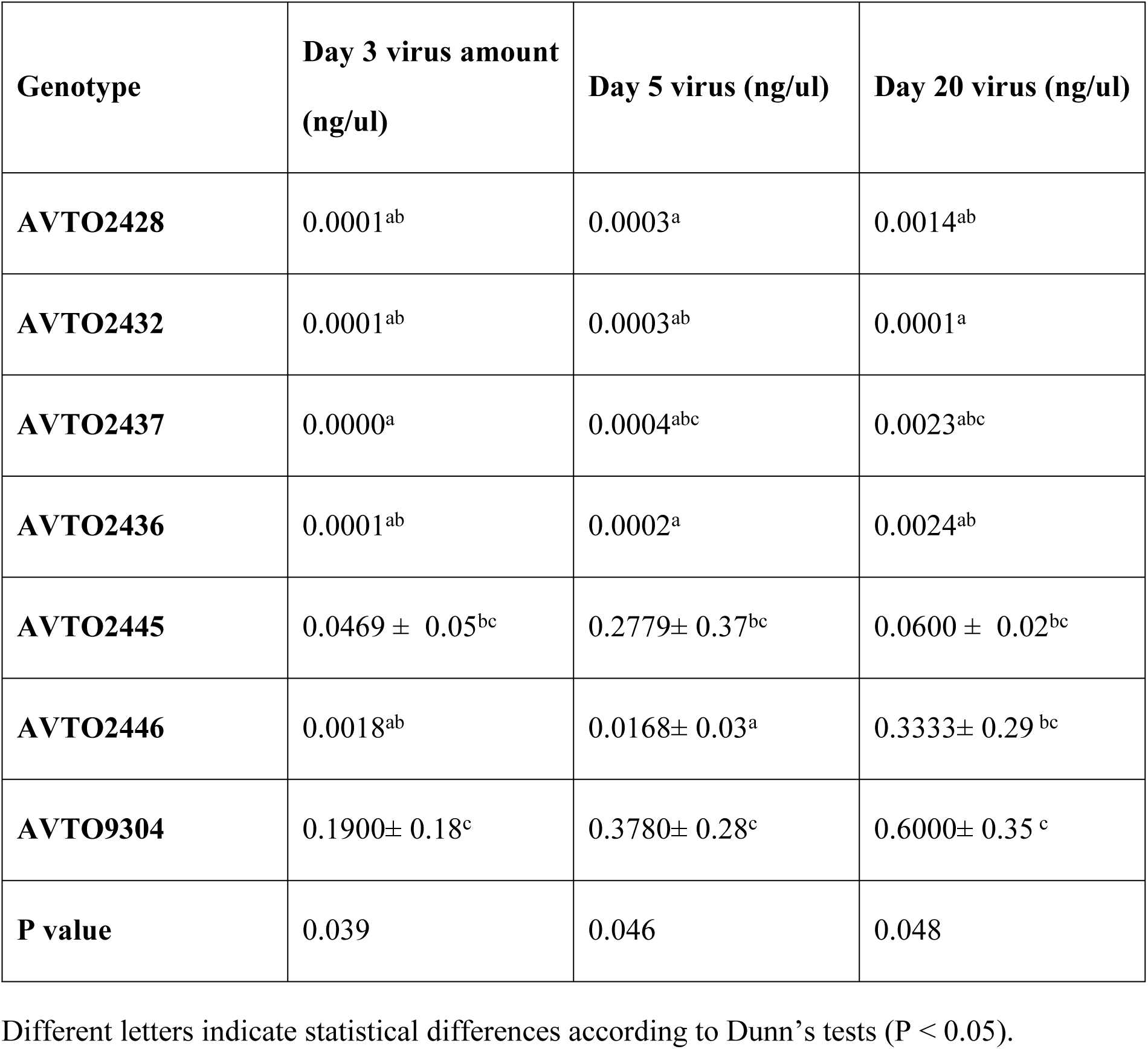
Virus accumulation over time in choice assay.

**Fig 6.**
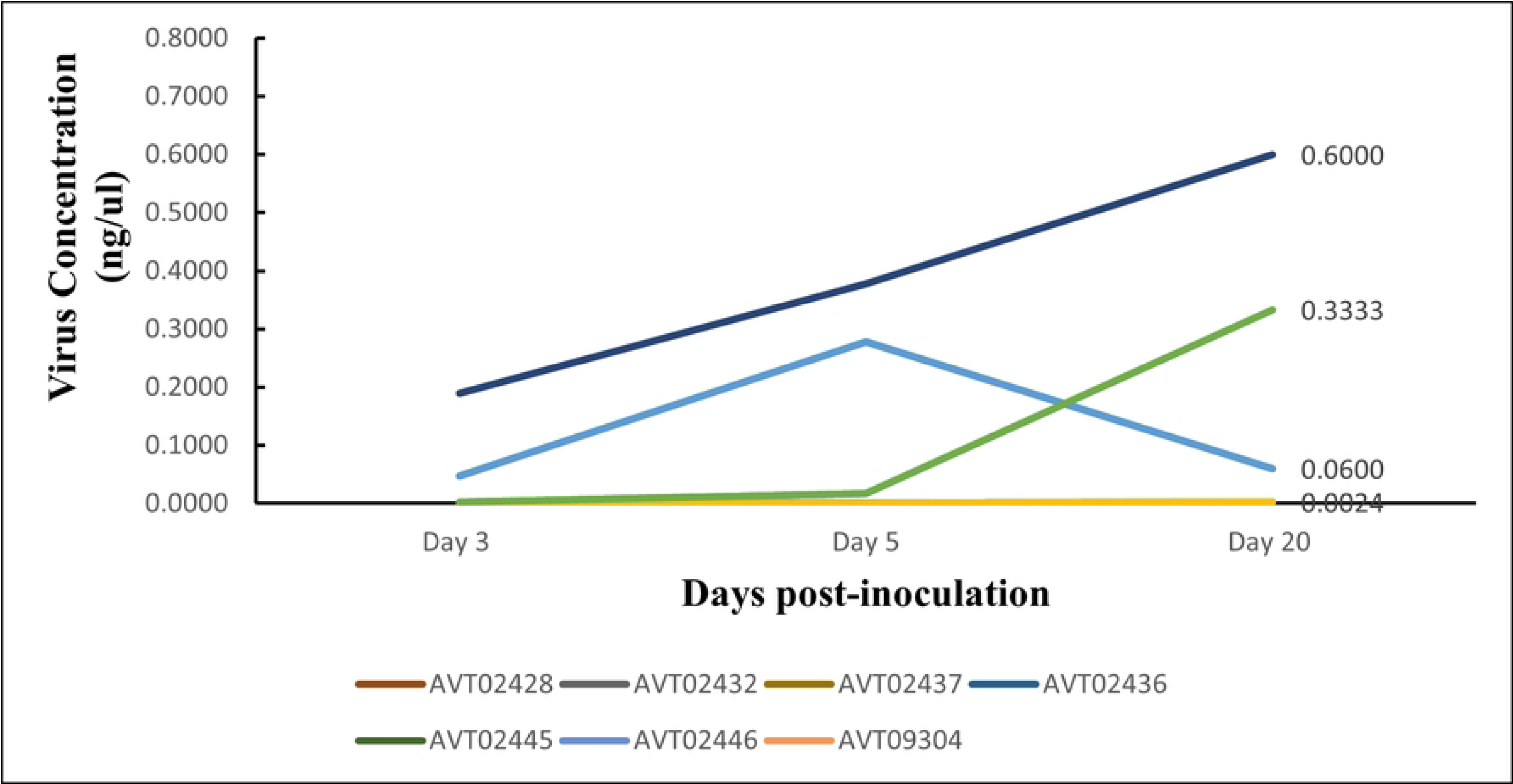
Virus accumulation over time.

### Disease scoring

The summary of the results of disease scoring is presented in Table 4 and Fig 7. The multi-disease and insect-resistant plants consistently showed lower disease score and no to very mild symptoms. At 35 dpi, the effectiveness of multi-disease and insect-resistant plants is demonstrated by an average severity rating of 1, indicating that all plants were healthy. In contrast, single-resistant plants and the susceptible check had average severity ratings of 2.5 and 4.44, respectively.

**Table 4.** Results of disease scoring.

| Genotype | Day 21 TY DSI | Day 28 TY DSI | Day 35 TY DSI |
| --- | --- | --- | --- |
| AVTO2428 | 1.00<sup>a</sup> | 1.00<sup>a</sup> | 1.00<sup>a</sup> |
| AVTO2432 | 1.00<sup>a</sup> | 1.00<sup>a</sup> | 1.00<sup>a</sup> |
| AVTO2437 | 1.11± 0.19<sup>ab</sup> | 1.00<sup>a</sup> | 1.00<sup>a</sup> |
| AVTO2436 | 1.00<sup>a</sup> | 1.00<sup>a</sup> | 1.00<sup>a</sup> |
| AVTO2445 | 2.00<sup>bc</sup> | 2.89± 0.19<sup>b</sup> | 2.67± 0.34<sup>b</sup> |
| AVTO2446 | 1.17± 0.29<sup>ab</sup> | 1.83± 0.29<sup>ab</sup> | 2.33± 0.76<sup>ab</sup> |
| AVTO9304 | 3.44± 0.2<sup>c</sup> | 4.44± 0.39<sup>b</sup> | 4.44± 0.39<sup>b</sup> |
| P value | 0.008 | 0.003 | 0.003 |

Different letters indicate statistical differences according to Dunn's tests ( $P < 0.05$ ).

**Fig 7.**
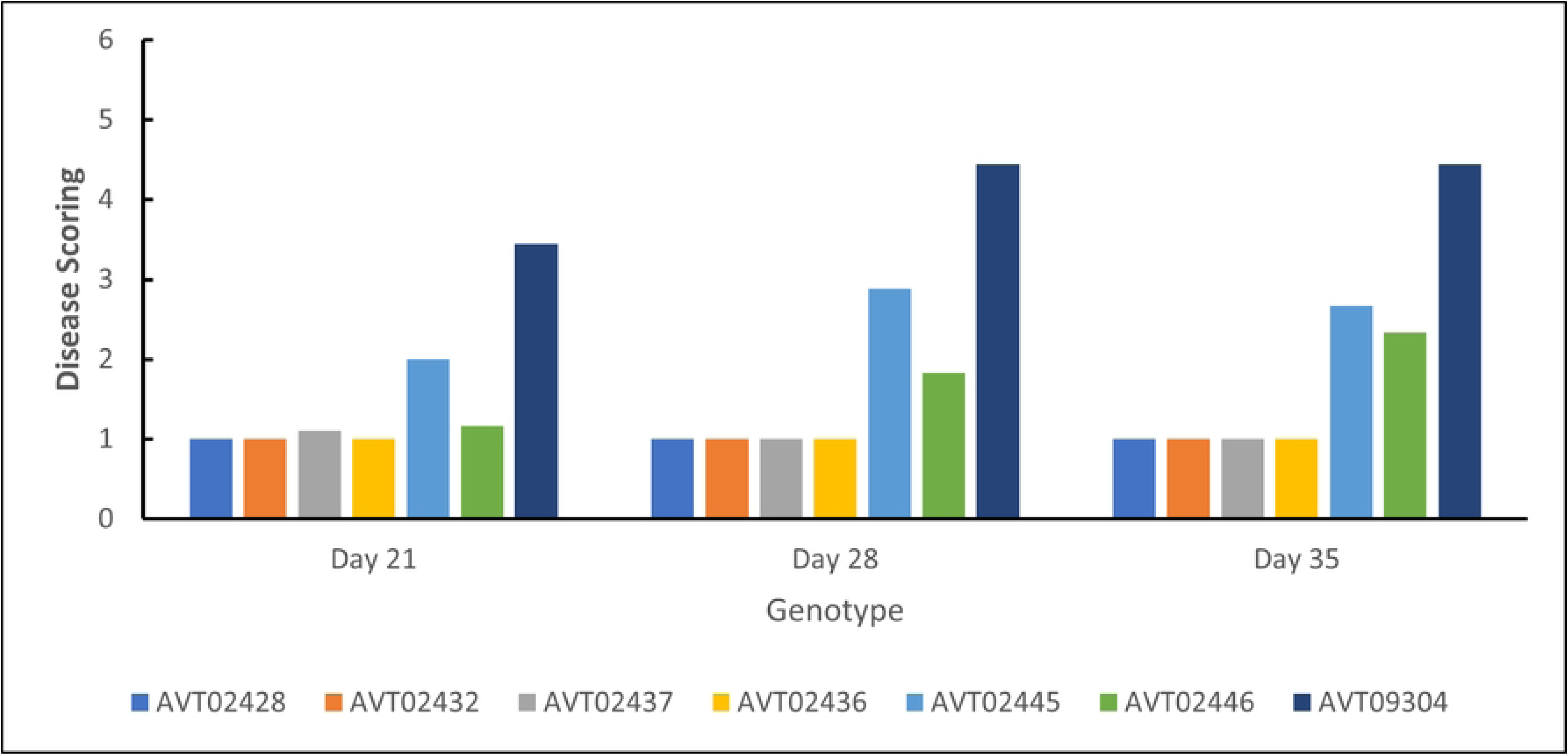
Results of disease scoring.

Kruskal-Wallis test indicated a significant difference in disease score across the 7 genotypes in both the assays. Post-hoc comparisons using Dunn’s test showed that at 35 dpi, all four lines differed significantly from *AVTO2445* and *AVTO9304*.

## Discussion

This study demonstrates that combining resistance to *B. tabaci* based on acylsugar secretion with TYLCV resistance genes (*Ty-1/Ty-3*) in multi-disease and insect-resistant tomato lines significantly reduced the accumulation of the TYLCV compared to plants carrying either only insect or TYLVC resistance, and, as expected, also compared to the susceptible check.

It is expected that the choice bioassay would reveal lines that confer both antixenosis and antibiosis resistance mechanisms [25]. The initial adult choice of a host plant for feeding is influenced more by preference factors. The preference assays confirm that the lines with high acylsugar secretion like *AVTO2428*, *AVTO2432*, *AVTO2437*, *AVTO2436,* and *AVTO2446* are less preferred, whereas *AVTO2445* and *AVTO9304* are susceptible. Previous studies have suggested that glandular trichomes on leaves and acylsugar secretion were associated with reduced attractiveness and resistance to *B. tabaci* [12,15–17,26–28]. Our results aligned these findings with the whitefly-resistant lines showing higher acylsugar concentration. The study revealed that high acylsugar levels were negatively correlated with preference by whitefly.

Whitefly feeding, besides vectoring viruses, causes a range of other disorders. The multi-disease and insect-resistant lines might help to reduce the irregular ripening disorder caused by *B. tabaci* on tomatoes by lowering whitefly infestations [17]. Host resistance to whiteflies presumably also reduces spread of other viruses transmitted by whiteflies, such as Tomato chlorosis virus [29], thereby lowering the risk of the emergence of recombinant viruses due to mixed infection. Due to the combination of different resistance mechanisms, multi-disease and insect-resistant lines might also exert lower selection pressure on the virus to evolve [30].

The resistance to adult *B. tabaci* in the tomato lines was associated with the high acylsugar concentration in these lines, which reduced the plant’s attractiveness to the insect and therefore reduced settling and oviposition. Rodríguez-López *et al.*(2011) found that the insect-resistant plants showing deterrence due to trichomes and acylsugar concentrations alter the feeding behaviour of the insect after it lands on a plant and affects virus acquisition by decreasing the ability to start probing. The tested whitefly resistance does not completely block TYLCV acquisition and transmission in vector-resistant plants. Hence, combining vector resistance with virus resistance to reduce disease development is necessary [17].

The graph of virus accumulation over time indicates that when the whiteflies are given a free choice between the genotypes, more whiteflies are attracted to the susceptible check *AVTO9304*, leading to high initial virus inoculation coupled with a lack of any virus resistance mechanism to reduce viral multiplication. The high initial virus concentration multiplies faster, leading to high virus accumulation. In the virus-resistant genotype *AVTO2445*, even though the initial virus concentration is high due to susceptibility to whiteflies, the virus multiplication is suppressed due to presence of *Ty-1/Ty-3* resistant genes. In the vector-resistant genotype *AVTO2446*, the initial virus concentration is low due to whitefly resistance, but due to absence of any mechanism to reduce viral multiplication, even a small initial viral load multiplies at a high rate, resulting in high virus accumulation over time and leading to disease development. In contrast, the multi-disease and insect-resistant lines consistently show lower virus concentrations. At 20 dpi, the multi-disease and insect-resistant line *AVTO2432* performed significantly better than the single-resistant plants and the susceptible check. This could be attributed to the combined resistance targeting the virus and its vector, creating a more comprehensive barrier against TYLCV infection and accumulation. These cultivars repel whitefly infestation and exhibit significant efficacy in impeding viral proliferation within the plant tissues.

Lower viral loads are directly correlated with reduced symptom severity, demonstrating the effectiveness of multi-disease and insect resistance in alleviating TYLCV symptoms. The mild symptoms observed in the dual-resistant plants can be attributed to their enhanced ability to restrict virus replication and movement within the plant tissues and reduce virus acquisition.

In conclusion, this study demonstrates that the multi-disease and insect-resistant tomato lines significantly reduce the accumulation of TYLCV and disease symptoms. The high acylsugar concentration reduces the preference of whiteflies for settling, resulting in reduced initial viral load. Together with TYLCV resistance this leads to lower virus accumulation over time and less severe symptoms, underscoring the effectiveness of these lines in managing TYLCV. These lines offer a robust and sustainable solution to prevent TYLCV damage in tomato breeding programs. These lines can reduce the reliance on chemical controls to combat disorders caused by whiteflies and lead to more sustainable agricultural practices. The study highlights the potential of multi-disease and resistant tomato lines as a key component of IPM strategies to manage whitefly-transmitted TYLCV effectively. These lines are a breakthrough in resistance breeding for TYLCD management. The promising results from this study pave the way for further research and development of multi-disease and insect-resistant lines to combat TYLCV and other whitefly-transmitted viral diseases.

Further research should focus on conducting field trials under diverse environmental conditions and with different TYLCV strains to validate the effectiveness of these lines in real-world agricultural settings, exploring the long-term durability of resistance and the potential for resistance breakdown. Further, the effect of pyramiding insect resistance with other TYLCV resistance genes and resistance genes for other viruses of tomato into these multi-disease and insect-resistant lines needs to be investigated to optimize the implementation of these resistant lines as an IPM component to maximize their benefits in agricultural practice and further improving the horticultural traits such as fruit size and yield according to the market demands for better adoption by farmers.

